# Optimized cell culture conditions promote *ex-vivo* manipulation and expansion of primitive hematopoietic stem cells for therapeutic gene editing

**DOI:** 10.1101/2022.01.11.475795

**Authors:** Rajeev Rai, Winston Vetharoy, Asma Naseem, Zohar Steinberg, Adrian J Thrasher, Giorgia Santilli, Alessia Cavazza

**Affiliations:** Infection, Immunity and Inflammation Research and Teaching Department, Great Ormond Street Institute of Child Health, University College London, 30 Guilford Street, London WC1N 1EH, UK

## Abstract

During the last few years, gene editing has emerged as a powerful tool for the therapeutic correction of monogenic diseases. CRISPR/Cas9 applied to hematopoietic stem and progenitor cells (HSPCs) has shown great promise in proof-of-principle preclinical studies to treat haematological disorders, and clinical trials using these tools are now underway. Nonetheless, there remain important challenges that need to be addressed, such as the efficiency of targeting primitive, long-term repopulating HSPCs and expand them *in vitro* for clinical purposes. Here we have tested the effect exerted by different culture media compositions on the ability of HSPCs to proliferate and undergo homology directed repair-mediated knock-in of a reporter gene, while preserving their stemness features during *ex-vivo* culture. We tested different combinations of compounds and demonstrated that by supplementing the culture media with inhibitors of histone deacetylases, and/or by fine-tuning its cytokine composition it is possible to achieve high levels of gene targeting in long-term repopulating HSPCs both *in vitro* and *in vivo*, with a beneficial balance between preservation of stemness and cell expansion, thus allowing to obtain a significant amount of edited, primitive HSPCs compared to established, state-of-the-art culture conditions. Overall, the implantation of this optimized ex vivo HSPC culture protocol will improve the efficacy, feasibility and applicability of gene editing and will likely provide one step further to unlock the full therapeutic potential of such powerful technology.

## Introduction

Over the past decade, there has been a global upsurge in the identification and discovery of novel gene editing (GE) tools, which has endowed the scientific community with the ability to artificially modify genetic information, unlocking the potential of traditional medicine to translate into new therapeutic approaches. In particular, the CRISPR/Cas9 system has proven to be a versatile platform for gene addition and deletion strategies in the arena of blood disorders and ongoing clinical trials to treat hemoglobinopathies are showing encouraging results.^1^ The CRISPR/Cas9 platform relies on a DNA-binding guide RNA (gRNA), which is complementary to the target DNA sequence, and a Cas9 endonuclease that creates a double strand break upon binding to the target site on the DNA, triggering the activation of two main endogenous repair pathways: nonhomologous end joining (NHEJ) and homology-directed repair (HDR). Each of these pathways could be exploited for therapeutic purposes. HDR mediates the accurate repair of the cut by utilizing a DNA sequence homologous to the region flanking the double strand brake as a template. As this process results in the insertion of a correct DNA sequence, this pathway can be harnessed when treating those diseases for which correcting or adding a genetic element may lead to a therapeutic benefit, mimicking the gene addition approach achieved through viral gene therapy.

In the context of blood disorders such as primary immunodeficiency diseases (PIDs), homology directed repair (HDR)-mediated editing is probably the most frequently used repair pathway to correct loss-of-function mutations underlying the disease, by either site-specific insertion of a gene or correction of patient-specific mutations.^2^ From the very first PID gene therapy clinical trial, successful treatment outcomes have been generated from the *ex vivo* modification of haematopoietic stem and progenitor cells (HSPCs). Indeed, long-term repopulating HSPCs (HSCs) are the ideal target for gene editing of various types of inherited haematological conditions. The therapeutic benefit of gene-corrected HSPCs depends on their capacity to engraft and provide long-term production of healthy blood lineage progenitors while maintaining renewable stem cells in transplanted patients.^3^

When applying genome editing with the aim to repair the functional and phenotypic defects of a disease, one not only needs to choose which specific tool to utilize, but also the delivery route, dose, and timing of editing reagents. This is to ensure that high HDR efficiency, and negligible cytotoxic and off-target effects are achieved in the cell type of interest. Overall, the application of genome editing to HSPCs has shown substantial benefits toward functional gene correction for different types of PIDs. Depending on the type and location of the mutation, researchers have shown that it is possible to fully recapitulate a physiological gene expression pattern and restore a functional immune response by HDR-mediated editing of the desired genomic locus in HSPCs. Evidence has also shown the ability of patient-derived edited cells to engraft the bone marrow and differentiate into various immune cells *in vivo*.^4-6^ In particular, our lab has recently demonstrated the successful development of a CRISPR/Cas9-based gene editing platform to treat Wiskott-Aldrich Syndrome (WAS) and showed the feasibility and the superiority of the gene editing strategy compared to viral vector-based gene therapy in correcting the WAS defects both *in vitro* and *in vivo*.^7^

Achieving sustained and higher levels of targeted integration is one of the ultimate goals to completely cure diseases by gene editing. However, following the *ex vivo* delivery of the editing reagents, NHEJ is the preferential pathway utilized to correct the DSB in nondividing cells, such as HSPCs. To overcome this issue, various groups have implemented strategies to either inhibit NHEJ,^8^ or increase the frequency of HDR,^9^ and have optimized timing and dosage of the editing reagents^6,10,11^ to enhance knock-in efficiency. The most successful approach so far has probably arisen from the attempt to promote HSC cell cycle progression to increase the engagement of HDR components and hence the knock-in exogenous sequences, which has been accomplished by the use of cell cycle modulators.^12-14^ Overall, the implementation of these methods to circumvent the limitations of the technology have led to outstanding levels of gene engineering in HSPCs *in vitro*, with >70% of correction rates achieved with GE by us and others in HSPCs,^4,5,7,10,14-17^ matching those frequently obtained by gene therapy gene addition strategies and predicted to be curative for the majority of blood disorders. However, while good engraftment rates have been observed following transplant of HSPCs electroporated with Cas9:gRNA RNP into immunodeficient mice, the persistence of edited cells in the hematopoietic tissues decreases significantly within 8–16 weeks after transplant and in serial transplantation experiments. This is an important and pressing issue for the gene therapy field, as inadequate engraftment and persistence of corrected cells hampers the broader application of these technologies to the treatment of blood disorders that require high chimerism post transplantation and that do not display strong selective advantage of corrected cells. The decline in the frequency of corrected cells *in vivo* could be due to the inefficient HDR-mediated editing in quiescent long-term repopulating HSCs, or their inability to self-renew upon their manipulation *in vitro*, including exposure to the editing reagents and culture conditions. Different strategies have been put in place in recent years to maintain and expand the primitive pool of self-renewing HSCs, such as the use of small molecules (UM171, PGE2, and StemRegenin1), as well as the optimization of culture conditions and timing of delivery of the editing reagents to HSPCs to preserve their engraftment potential.^6,10,11,14^

In this study, we aim to identify HSPCs culture conditions that support targeting of primitive HSCs while favouring their expansion *in vitro* and transplantation *in vivo*. We show that the addition of IL-3 in the stem cell medium greatly supports HSPCs expansion and HDR-mediated transgene knock-in while promoting cell differentiation and limited maintenance of long-term repopulating HSCs. At the contrary, media containing IL-6 better preserves HSPC stemness at the expense of cell expansion and HDR frequencies. We therefore tested different combinations of compounds and demonstrated that by supplementing the culture media with inhibitors of histone deacetylases, and/or by fine-tuning its cytokine composition it is possible to achieve high levels of gene targeting with a beneficial balance between preservation of stemness and cell expansion.

## Results and discussion

To assess the impact of media composition on HSPCs expansion, preservation of stemness and HDR efficiency, we utilized a CRISPR/Cas9 platform we previously designed for the *WAS* locus, comprised of a gRNA targeting the start codon of *WAS*, a HiFi Cas9 and an AAV6 donor vector carrying a GFP reporter cassette flanked by 400-bp homology arms.^7^ This platform has been chosen for its optimal performance, with consistent high levels of gene knock-in across different CD34+ HSPC donors and experimental replicates and good rates of engraftment of edited cells *in vivo*. We first focussed our attention on the basal composition of the medium used to culture HSPCs with the purpose of gene manipulation. Traditionally, HSPCs *ex vivo* manufacturing protocols provide the use of a serum-free culture media supplemented with a predefined stem cell cocktail that includes Fms-related tyrosine kinase 3 (FLT3), thrombopoietin (TPO) and stem cell factor (SCF), with the addition of basal cytokines to promote cell fitness and proliferation. We first tested the impact of IL-3 or IL-6, the two most commonly used cytokines for HSPC culture, on HSPCs’ ability to proliferate and undergo HDR-mediated knock-in of a PGK-GFP reporter cassette into the *WAS* locus. To this aim, HSPCs harvested from the peripheral blood (PB) of more than 3 different healthy donors were cultured for two days in a basal, FLT3-TPO-SCF-containing media supplemented with either IL-3 or IL-6 and then electroporated with a Cas9-gRNA ribonucleoprotein complex (RNP) and transduced with the AAV6 donor vector using established protocols;^7^ cells were kept in culture for two additional days, after which they were harvested to assess gene editing efficiency and cell phenotype. HSPCs cultured in IL-3 containing media showed comparable levels of NHEJ-mediated repair of the double strand break induced at the *WAS* locus compared to IL-6 (**Figure 1A**), with however a higher frequency of HDR-mediated GFP integration (average 42% and 27%, respectively; **Figure 1B**). As the engagement of the HDR machinery is strictly dependent on cell proliferation, we assumed that the increased HDR frequency was a direct consequence of increased HSPC cycling mediated by IL-3; indeed, when measuring cell numbers after 6 days of culture we noticed a >2-fold increase in total cell numbers when adding IL-3 *versus* IL-6 (**Figure 1C**), which, combined with higher knock-in rates, translated into an average of 3-fold more HSPCs with a reporter cassette correctly knocked-in (**Figure 1D**). To investigate the stem cell phenotype of HSPCs cultured in the two different media, we sought to determine the frequency of primitive, long-term repopulating HSCs and multipotent progenitors (MPPs) in the bulk HSPC populations cultured in two different media by flow cytometry using a well-established panel of markers.^18^ We found a striking reduction of HSCs in the bulk of CD34+ HSPCs cultured in the presence of IL-3 *versus* IL-6, while MPPs were better preserved by IL-3 compared to IL-6, at both 4 and 7 days after editing (**Figure 1E,F**). Our data are in line with previous studies and observations, where IL-3 has been reported to promote the proliferation of HSPCs, together with increased cell differentiation and reduction of their repopulating potential *in vivo*^19^, while IL-6, SR and UM171 containing media have been shown to maintain high frequency of HSCs *in vitro* and partially *in vivo*.^6,10^ Our results highlighted important features of both media used in the context of gene editing. As mentioned earlier, proliferation is a key aspect for efficient gene correction by HDR and so is the availability of high numbers of edited HSCs to be transplanted into patients; indeed one critical limiting factor to the successful clinical translation of gene editing is the relatively low numbers of HDR-corrected HSCs obtained during the manufacturing process, which may lead to graft failure and lack of therapeutic benefit when infused into patients, especially those suffering from conditions that require high levels of chimerism to be treated. In light of these considerations, a medium containing IL-3 could be favourably used for gene editing of HSPCs to obtained high numbers of corrected cells. However, the pronounced cell differentiation induced by IL-3 is likely to influence the long-term bone marrow repopulating ability of edited HSPCs, resulting in the loss of engrafted and corrected cells once transplanted *in vivo*. We reasoned that optimising culture conditions by integrating the advantages of the two media could lead to an optimal balance between proliferation and stem cell preservation; therefore new media configurations were conceived with the aim to either 1) increase the stemness preserving capacity of IL-3 containing media (medium A); or 2) increase the proliferation ability of the IL-6 containing media (medium B). To these aims, we investigated the effect of supplementing IL-3 media with HDAC inhibitors (HDACi) (medium C), which were previously shown to successfully promote a HSC phenotype through the regulation of epigenetic plasticity and chromatin structures which are critical for the maintenance of the primitive status of HSCs.^20-22^ In parallel, we also tested the addition of IL-3 to the IL-6, UM171 and SR-containing media (medium D).

**Figure 1.**
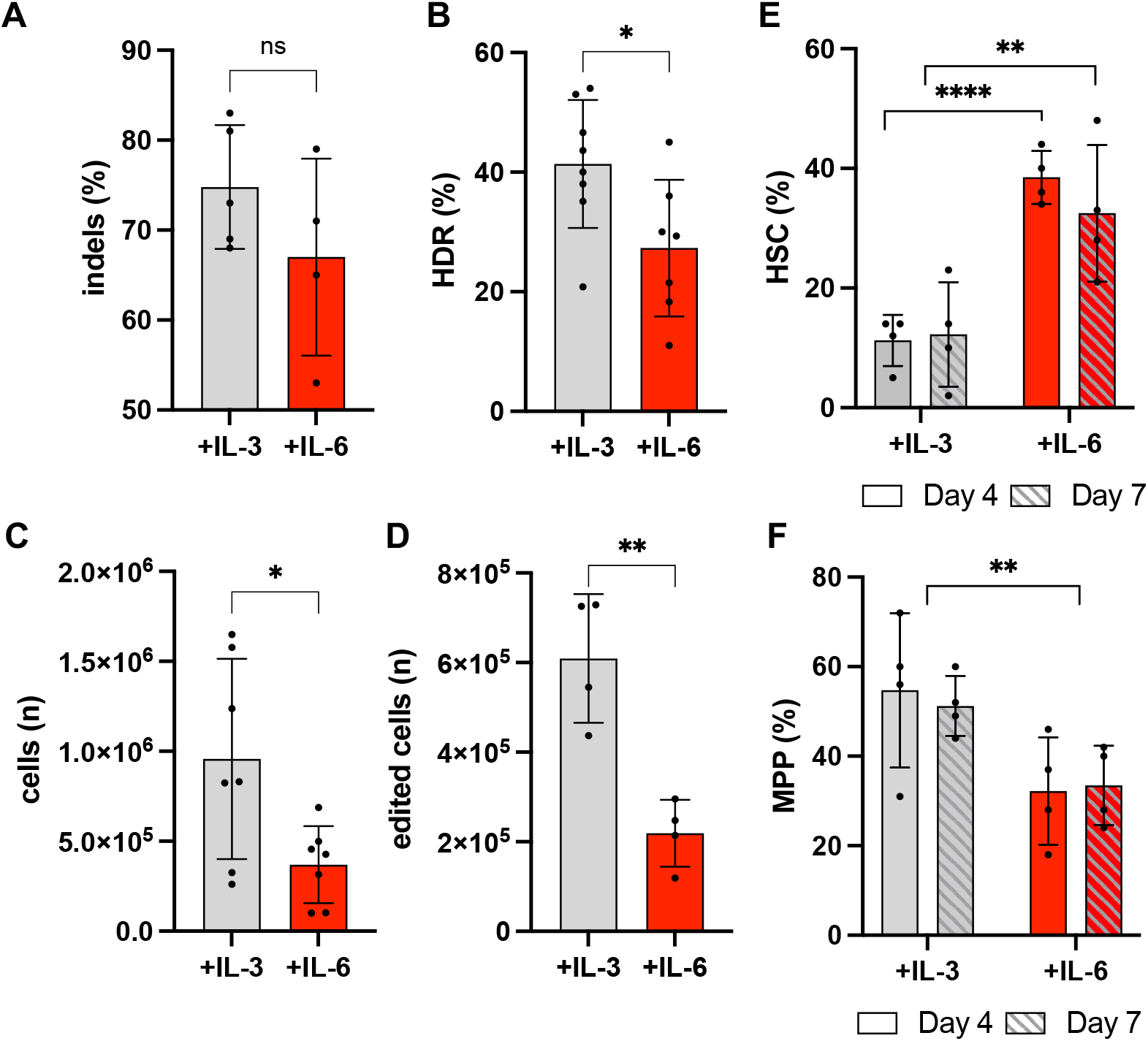
Comparison of *ex vivo* culture and manipulation of HSPCs grown in media supplemented with either IL-3 or IL-6. Frequency of **A)** NHEJ-mediates repair (indels) and **B)** HDR-mediated knock-in of a PGK GFP reporter cassette at the *WAS* locus in CD34+ HSPCs cultured in a IL-3 or IL-6 supplemented stem cell medium; **C)** Total number of cells and of **D)** edited cells retrieved in either medium after 6 days of culture; **E)** frequency of HSCs and **F)** MPPs detected in the CD34+ bulk cultured in either medium 4 and 7 days after gene editing (6 and 9 days of culture, respectively). Data in Figure 1 are presented as mean ± SD, with n=4 biological replicates. *P*-values were calculated using one-way ANOVA with Tukey’s comparison test (e,f) or two-tailed unpaired Student’s *t* test (a-d).

To accurately estimate rates of gene editing and preservation of stem cell phenotypes in different subpopulations of stem and progenitor cells, HSCs, multipotent progenitors (MPPs) and CD38+ committed progenitors were sorted from PB-derived CD34+ HSPCs immediately after cell thawing and placed in the four different culture media under investigation. Two days post sorting, the three populations, as well as unsorted CD34^+^ cells, were edited by delivery of the Cas9/gRNA RNP and the AAV-PGK-GFP donor vector and rates of targeted integration, as well as the frequency of HSCs and MPPs in the culture, were determined 4 days after editing, for a total of 6 days of cell culture (**Figure 2A**). This timeline was purposely chosen to mimic a potential protocol of HSPC gene therapy manufacturing, which is usually 4-6 days long.

**Figure 2.**
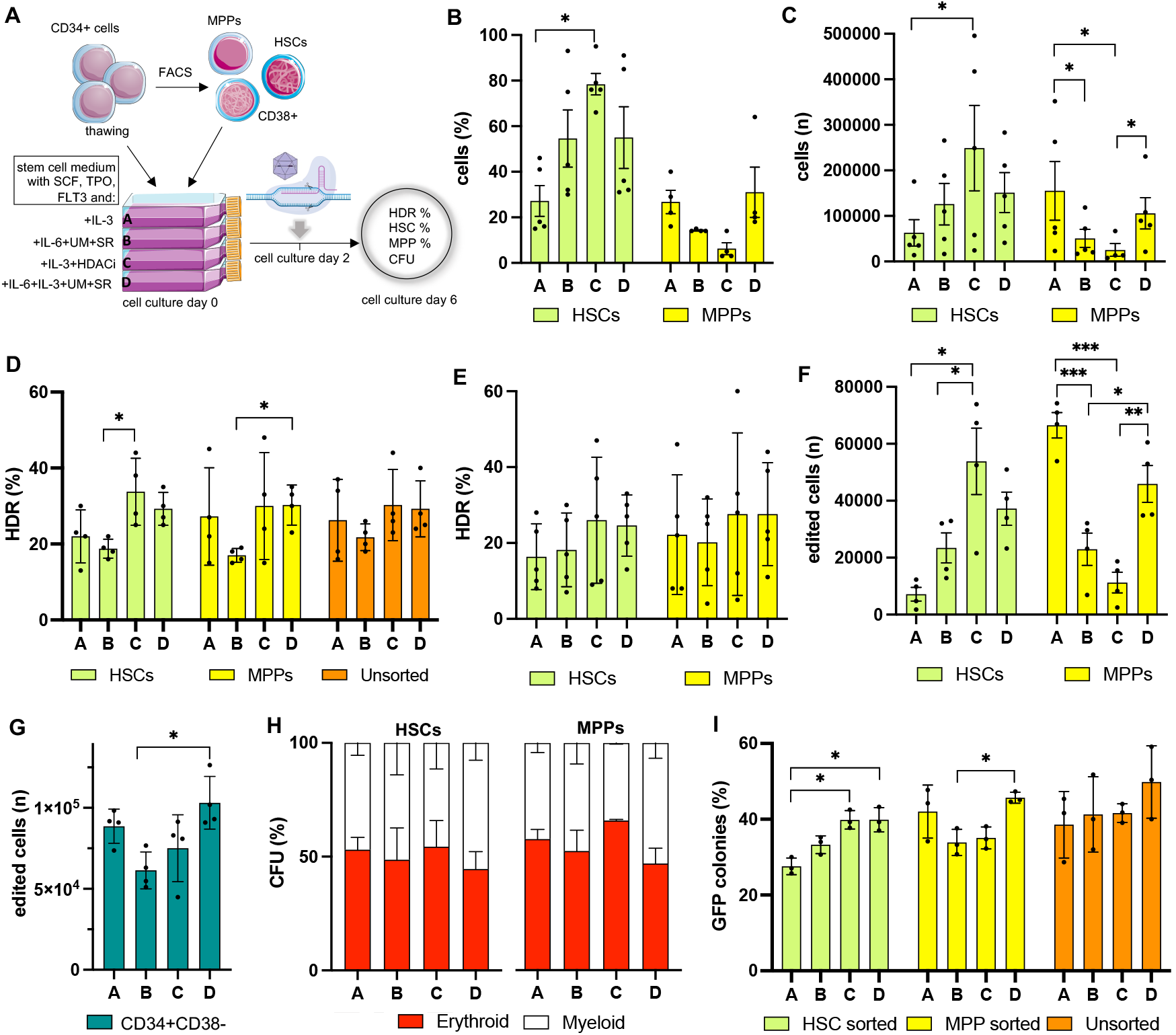
Evaluating the frequency of gene knock-in and preservation of stemness in sorted populations. **A)** Schematic representation of the experimental procedure. CD34+ cells are thawed and FACS sorted into primitive HSCs and MPPs. HSCs, MPPs and unsorted cells are then culture into four different media (A-D) containing different combinations of cytokines and stem cell factors. After 2 days of culture, sorted populations are gene edited with the CRISPR/Cas9 platform and the AAV6 donor vector containing a PGK-GFP reporter cassette. After 4 additional days of culture, cells are harvested and analysed. **B)** Percentage and **C)** number of cells in the HSC or MPP sorted populations that retain an HSC or MPP phenotype after gene editing and 6 days of culture in different media; **D)** Percentage and **F)** number of cells harbouring a GFP reporter cassette in the *WAS* locus in the sorted populations cultured in the different media; **G)** number of edited repopulating CD34+ CD38-cells in each medium; **H)** Plots representing the percentage of myeloid (white) and erythroid (red) colonies formed in methylcellulose by sorted and edited HSC and MPP populations cultured in the different media; **I)** Plots representing the percentage of GFP-positive colonies formed in methylcellulose by sorted and edited HSC and MPP populations or unsorted and edited CD34+ HSPCs cultured in the different media. Data in Figure 2 are presented as mean ± SD, with n=4 biological replicates in all panels except for H and I where n=3. *P*-values were calculated using one-way ANOVA with Tukey’s comparison test.

At day 6 of cell culture, medium C showed the highest preservation of HSCs (average 79% of HSCs in the HSC sorted population), with a 3-fold increase compared to IL-3 only containing medium (medium A), highlighting the ability of HDACi to preserve HSCs in culture. Given that the post culture counts of cells for medium C were equal or lower to those observed for the other media (**Supplementary Figure**), we speculate that the enhancement in the HSC fraction mediated by medium C could be ascribed to stem cell preservation rather than their expansion. Cells cultured in medium B and D, IL-6 containing media, also mantained a sensible and comparable fraction of HSCs (average 57%), while sorted cells grown in medium A showed the most pronounced loss of stemness, confirming the results obtained previously. The HSC preserving effect exerted by medium C was specific to the primitive stem cell pool, as no significant differences with the other media were observed in the MPP fraction (**Figure 2B**); however, we could notice a favourable trend for media A and D in the preservation of MPPs after 6 days of culture. When looking at the total number of cells still showing a primitive phenotype in HSC sorted cell populations at day 6 of culture (“preserved” HSCs), medium C outperformed all the other media and yielded 3 times more HSCs than medium A (**Figure 2C**). Again, this effect was specific to the stem cell population, as the medium was found to yield the lowest number of MPPs in the MPP sorted sample, while medium A and D were consistently very effective in the preservation of MPPs. HDR-mediated knock-in of the reporter cassette was detected in all cell populations and all media, with an overall comparable amount of GFP-expressing cells; Medium C showed the highest levels of editing in HSCs followed by Medium D, the latter being the best supporter of editing in MPPs as well (**Figure 2D**). No major differences in the frequency of knock-in could be evidenced in the unsorted population, highlighting the importance of studying the effects of culture conditions and HDR efficiency in carefully selected populations rather than retrospectively in the CD34+ bulk. Medium C yielded the highest number of edited cells displaying an HSC phenotype within the HSC-sorted population at culture day 6 (**Figure 2E**), while both Medium A and Medium D yielded the highest number of edited and preserved MPPs. Overall, when looking at the number of cells within the CD34+ CD38-population, which contains both HSCs and MPPs and has been shown to be the major contributor of long-term engraftment in immunodeficient mice when transplanting edited HSPCs,^13,23^ medium D outplays all the other conditions, providing the highest number of edited, bone marrow repopulating cells (**Figure 2G**). Sorted and edited HSCs and MPPs and unsorted CD34^+^ cells (**Figure 2H** and **Supplementary Figure**) showed similar clonogenic potential in CFU assays, without any significant lineage skewing compared to controls and among media. HSC-sorted cells cultured in medium C and D yielded a higher output of GFP^+^ colonies in methylcellulose, in line with the evidence of superior amount of cells preserving an HSC phenotype in optimized culture conditions and higher editing rates (**Figure 2I**); the high proportion of GFP+ colonies in the medium D condition observed in both MPPs and unsorted samples matches the previous findings of increased phenotype preservation and gene editing frequencies in HSCs and MPPs cultured in the presence of both IL-3 and IL-6.

We next evaluated whether HSPCs cultured and manipulated in the four different media retain their capacity to repopulate the bone marrow and differentiate into all the hematopoietic lineages. CD34+ HSPCs from two different healthy donors were thawed, placed in culture in media A-D and edited following our standard protocol; at day 4 of culture, cells were transplanted into 8-weeks-old sub-lethally irradiated immunodeficient non-obese diabetic (NOD)-SCID Il2rg^−/−^ (NSG) mice together with uncultured HSPCs as a control (**Figure 3A**). A targeted integration frequency of 30-40% was detected in all the samples *in vitro*, with no significant differences among the media used (**Figure 3B**). Short-term human engraftment was measured in the PB of mice at week 8 after transplant, with the highest chimerism observed in mice transplanted with unmanipulated, uncultured HSPCs, followed by those transplanted with cells cultured in medium D (**Figure 3C**). The same trend was visible at 14 weeks post-transplant in the bone marrow (BM) of mice, with Medium D showing a strikingly higher frequency of engraftment compared to the other experimental groups and, importantly, with similar levels compared to unmanipulated controls (**Figure 3D**). We investigated the composition of the CD45+ engrafted human population in the BM of transplanted mice and observed a higher number of CD38-CD90+ HSCs in the medium D group compared to the others, with a comparable quantity of cells as unmanipulated controls (**Figure 3E**); the amount of MPPs was similar between medium B and D, and the highest compared to the other conditions (**Figure 3F**). Flow cytometry analysis of T-cells (CD3+), myeloid progenitors (CD33), and B-cells (CD19+) showed no differences in lineage composition in the BM and PB among the experimental groups (**Supplementary Figure**), indicating correct differentiation of manipulated HSPCs.

**Figure 3.**
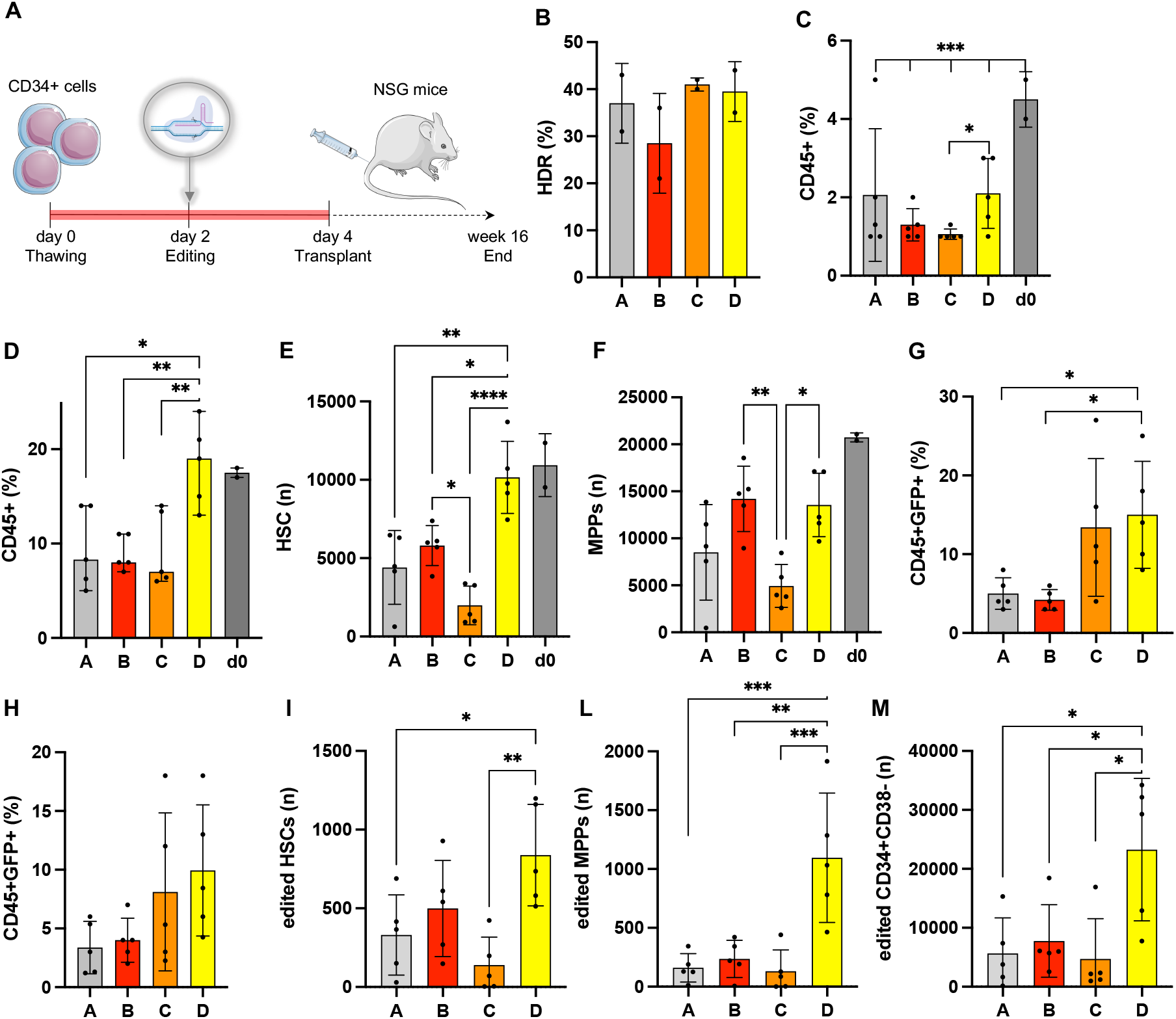
Evaluating the frequency of gene knock-in and in vivo hematopoietic reconstitution by gene edited HSPC. **A)** Schematic representation of the experimental procedure. CD34+ HSPCs from two different healthy donors are thawed, culture in media A-D and gene edited at day 2. Two days later, cells are harvested in transplanted into sub lethally irradiated NSG mice. 14-16 weeks after transplant, the experiment is terminated and the mice analysed to detect human cell engraftment. B) Rates of targeted integration achieved *in vitro* in HSPCs pre-transplant; C) Engraftment of human cells (CD45+) in the peripheral blood of NSG mice at 8 weeks post-transplant and D) in the bone marrow at 16 weeks post-transplant; E) number of human HSCs and F) MPPs detected in the bone marrow of transplanted mice; G) percentage of human cells with a GFP knocked in the WAS locus (CD45+ GFP+) in the bone marrow or H) peripheral blood of experimental mice at 16 and 8 weeks post-transplant, respectively; I) amount of edited HSCs and L)MPPs and M) CD34+ CD38-repopulating HSPCs found in the bone marrow of transplanted mice at 16 weeks post-transplant. Data in Figure 3 are presented as mean ± SD, with n=5 biological replicates in all panels except for A where n=2. *P*-values were calculated using one-way ANOVA with Tukey’s comparison test.

When examining the rate of targeted integration in human engrafted CD45+ cells in the BM (**Figure 3G**) and in the PB (**Figure 3H**) of experimental mice, an average of 12-15% of GFP expressing cells was detected in the Medium C and D groups; while we still observe a drop in the HDR rate *in vivo* compared to the *in vitro* counterpart, the reduction is much less evident when using the two best performing media, which show >2-fold increased frequency of engraftment compared to IL-3 only or IL-6 only media. Mice belonging to the Medium D group also present with the greatest amount of edited human HSCs and MPPs engrafted in the BM, and an overall significant increase in the number of edited CD34+CD38-repopulating cells (**Figure 3I-M**).

The data presented here identify the need of a fine-tuned balance between cues that promote cell proliferation, and hence HDR-mediated gene targeting, and preservation of stemness and repopulating potential when manipulating HSPCs *ex vivo* for therapeutic purposes. We showed that induction of proliferation and differentiation mediated by IL-3, which is beneficial for cell expansion and gene targeting, can be counterbalanced by the addition of stem cell preserving compounds, such as HDACi, IL-6, SR and UM171. Exposing HSPCs to HDACi seems to promote the maintenance of a primitive stem cell phenotype *in vitro*, however this effect is less evident when cells are transplanted *in vivo*. While the frequency of engrafted CD45+ cells and primitive HSCs in the BM of mice when using this medium is the lowest observed among the experimental conditions, the percentage of edited cells in the BM and PB of mice is among the highest. We speculate that the effect of the HDACi compound is to promote the quiescence of HSCs once engrafted, as already observed when culturing the cells in media supplemented with other compounds;^13^ however this observation must be confirmed by further studies on the effects exerted by HDACi on HSPCs. The addition of both of IL-3 and IL-6 to the HSPC medium is the most compelling strategy tested here as it combines the best of both worlds. Indeed, IL-3 contained in medium D is able to promote cell proliferation at sufficient levels to achieve HDR-mediated knock-in of the reporter cassette and cell expansion. On the other side, the presence of IL-6 and stem cell factors likely counteract the differentiation signals promoted by IL-3 and support the preservation of HSCs in culture at levels comparable to the current state-of-the-art media utilized in pre-clinical HSPC gene editing studies. The combination of these effects resulted in a higher amount of both HSCs and MPPs that are correctly gene edited, which also translates into the highest amount of GFP+ CD34+CD38-repopulating HSPCs among the conditions tested. It is worth mentioning that for a successful gene therapy application, both HSCs and MPPs must be corrected and infused into the patient, as the latter provides for a fast, short-term hemopoiesis within the first months after the transplant, ensuring a quick immune reconstitution to the patient.^24,25^ Hence, culturing HSPCs into a medium that promotes maintenance and editing of both populations represents an undisputed advantage for the success of the therapeutic intervention.

## Methods

### Human CD34^+^ HSPCs culture

Mobilised peripheral blood from healthy donors were isolated under written informed consent. Within 24 h of scheduled apheresis, CD34^***+***^ haematopoietic stem and progenitor cells (HSPCs) were purified using CD34+ Microbead kit (Miltenyi Biotec, UK) according to the manufacturer protocol. Purity was assessed by FACS staining with anti-human CD34 BV 421 antibody (clone 561, BioLegend, USA). For long-term storage, the cells were frozen in CryoStor cell cryopreservation media (Sigma-Aldrich). After thawing cells were cultured in StemSpan ACF (StemCell Technologies, USA) supplemented with SCF (100 ng/ml, Peprotech, UK), TPO (100 ng/ml, Peprotech), FLT3-ligand (100 ng/ml, Peprotech), IL-3 or IL-6 (60 ng/ml, Peprotech, UK). Cells were incubated at 37 °C/5% CO_2_ for 2 days prior to electroporation.

### Electroporation and transduction

After 2 days in culture, CD34^+^ HSPCs were electroporated using Neon Transfection kit (ThermoFisher Scientific). Briefly, 0.2 × 10^6^ cells were centrifuged and re-suspended in ribonucleoprotein (RNP) complex. The RNP was made by incubating gRNA and High Fidelity Cas9 protein (Integrative DNA Technologies, USA) at a molar ratio of 1:2 at 37 °C for 15 min. The condition for electroporation was 1600V, 10 ms, three pulses. Following electroporation, cells were seeded at concentration of 1 × 10^6^ cells per ml and incubated at 37 °C for 15 min after which AAV6 at 50,000 MOI (Vector genomes/cell) was added and incubated.

### Editing of CD34^+^ HSPCs subsets

At day 0, CD34^+^ HSPCs were thawed and stained with the following antibody panel: CD34 BV 421, CD38 APC-Cy7 (clone HIT2, BioLegend), CD90 PE-Cy7 (clone 5E10, BioLegend) and CD45RA APC (clone HI100, BioLegend). The cells were then FACS-sorted into different subsets including CD38+ cells, haematopoietic stem cells (HSCs) and multipotent progenitors (MPPs). Unsorted cells were taken as an experimental control. After 2 days, both sorted and unsorted cells were electroporated and transduced with AAV6_PGK-GFP. At days 5 and 12 post editing, the cells were stained with the above antibody panels to determine the percentage of HSCs and MPPs as well as evaluate GFP expression by FACS. Cell yields were calculated by adding 10 microliters of Count Bright Absolute Counting Beads (ThermoFisher Scientific), following the manufacturer’s protocol.

### Methylcellulose CFU assay

The colony-forming unit (CFU) assay was performed by seeding 500 cells in six-well plates containing MethoCult Enriched (StemCell Technologies) after 4 days of editing. After 14 days of incubation at 37 °C/5% CO_2_, different types of colonies including CFU-Erythroid (E), CFU-Macrophage (M), CFU-Granulocyte (G), CFU-GM and CFU-GEM were counted based on their morphological appearance.

### Transplantation of genome edited CD34^+^ HSPC in NSG mice

NSG adult female mice (6–8 weeks old) were purchased from Charles River and were sub-lethally irradiated (3 Gy) 24 h prior to transplantation. CD34^+^ HSPCs were thawed and cultured. At day 2 after editing, 0.5 × 10^6^ viable cells were injected via tail vein into mice with a 27 gauge × 0.5 inch needle. After 8 and 14 weeks post transplantation, mice peripheral blood from tail vein was lysed with 1× RBC lysis buffer (ThermoFisher Scientific) and stained with anti-human CD45 APC antibody (clone HI30, BioLegend) to evaluate human CD45+ engraftment by FACS. At week 14-16 weeks post transplantation, level of human engraftment and lineage composition were determined in the bone marrow and peripheral blood. Briefly, bone marrow cells were harvested by flushing tibiae and femurs with 1× PBS and passing through 40 μm strainer. Mononuclear cells were blocked with Fc blocking solution (BioLegend) and stained with the following antibody panels; CD45 BV421, CD19 PerCp Cy5.5 (clone HIB19, BioLegend), CD33 FITC (clone P67.6, BD Bioscience) and CD3 PE (clone OKT3, BioLegend) for FACS analysis. To determine human stem cell composition within bone marrow, cells were stained with the following antibody panels; CD45 BV421, CD34 FITC, CD38 APC-Cy7, CD90 PE-Cy7 and CD45RA PerCP Cy5.5.

### FACS analysis

BD FACSAria II (BD Bioscience) instrument was used for cell sorting of CD34^+^ HSPCs. For all flow cytometry analysis, a BD LSRII instrument (BD Bioscience) was used. Antibody dilutions used for FACS staining was 1 in 100. For data anlaysis, FlowJo v10 software (FlowJo LLC, USA) was used.

### Ethics and animal approval statement

For usage of human CD34^+^ HSPC from healthy donors, informed written consent was obtained in accordance with the Declaration of Helsinki and ethical approval from the Great Ormond Street Hospital for Children NHS Foundation Trust and the Institute of Child Health Research Ethics (08/H0713/87).

For experiments involving animals, mice were housed in a 12-h day-night cycle with controlled temperature and humidity. The ventilated cages had sterile bedding and everyday supply of sterile food and water in the animal barrier facility at University College London. Mice were bred and maintained in accordance with UK Home Office regulations, and experiments were conducted after approval by the University College London Animal Welfare and Ethical Review Body (project license 70/8241).

## Acknowledgements

We thank Dr. Ayad Eddaoudi (Flow Cytometry Core Facility, University College London) for assistance with flow cytometry and cell sorting; R.R., W.V., G.S. and A.J.T. were supported by the Wellcome Trust (104807/Z/14/ Z) and the NIHR Biomedical Research Centre at Great Ormond Street Hospital for Children NHS Foundation Trust and University College London. A.C., A.N. and Z.S. were supported by the GOSH Children’s Charity-LifeArc Therapeutic Accelerator Grant, the University College London Therapeutic Acceleration Support fund and the NIHR Biomedical Research Centre at Great Ormond Street Hospital for Children NHS Foundation Trust and University College London.

## Author contributions

R.R., W.V., G.T., Z.S. performed experiments, analysed data and reviewed the manuscript; G.S. and A.J.T. helped with study design and reviewed the manuscript; A.C. initiated the study, designed experiments, illustrated data, and wrote the manuscript. Competing interests A.J.T is on the Scientific Advisory Board of Orchard Therapeutics and Rocket Pharmaceuticals. The other authors declare no competing interests.

## References

1 Frangoul, H., Ho, T. W. & Corbacioglu, S. CRISPR-Cas9 Gene Editing for Sickle Cell Disease and beta-Thalassemia. Reply. N Engl J Med 384, e91, doi:10.1056/NEJMc2103481 (2021).

2 Rai, R., Thrasher, A. J. & Cavazza, A. Gene Editing for the Treatment of Primary Immunodeficiency Diseases. Hum Gene Ther 32, 43–51, doi:10.1089/hum.2020.185 (2021).

3 Ferrari, G., Thrasher, A. J. & Aiuti, A. Gene therapy using haematopoietic stem and progenitor cells. Nat Rev Genet 22, 216–234, doi:10.1038/s41576-020-00298-5 (2021).

4 De Ravin, S. S. et al. CRISPR-Cas9 gene repair of hematopoietic stem cells from patients with X-linked chronic granulomatous disease. Sci Transl Med 9, doi:10.1126/scitranslmed.aah3480 (2017).

5 Dever, D. P. et al. CRISPR/Cas9 beta-globin gene targeting in human haematopoietic stem cells. Nature 539, 384–389, doi:10.1038/nature20134 (2016).

6 Schiroli, G. et al. Preclinical modeling highlights the therapeutic potential of hematopoietic stem cell gene editing for correction of SCID-X1. Sci Transl Med 9, doi:10.1126/scitranslmed.aan0820 (2017).

7 Rai, R. et al. Targeted gene correction of human hematopoietic stem cells for the treatment of Wiskott - Aldrich Syndrome. Nat Commun 11, 4034, doi:10.1038/s41467-020-17626-2 (2020).

8 Maruyama, T. et al. Increasing the efficiency of precise genome editing with CRISPR-Cas9 by inhibition of nonhomologous end joining. Nat Biotechnol 33, 538–542, doi:10.1038/nbt.3190 (2015).

9 Song, J. et al. RS-1 enhances CRISPR/Cas9- and TALEN-mediated knock-in efficiency. Nat Commun 7, 10548, doi:10.1038/ncomms10548 (2016).

10 Charlesworth, C. T. et al. Priming Human Repopulating Hematopoietic Stem and Progenitor Cells for Cas9/sgRNA Gene Targeting. Mol Ther Nucleic Acids 12, 89–104, doi:10.1016/j.omtn.2018.04.017 (2018).

11 Genovese, P. et al. Targeted genome editing in human repopulating haematopoietic stem cells. Nature 510, 235–240, doi:10.1038/nature13420 (2014).

12 Kim, S., Kim, D., Cho, S. W., Kim, J. & Kim, J. S. Highly efficient RNA-guided genome editing in human cells via delivery of purified Cas9 ribonucleoproteins. Genome Res 24, 1012–1019, doi:10.1101/gr.171322.113 (2014).

13 Shin, J. J. et al. Controlled Cycling and Quiescence Enables Efficient HDR in Engraftment-Enriched Adult Hematopoietic Stem and Progenitor Cells. Cell Rep 32, 108093, doi:10.1016/j.celrep.2020.108093 (2020).

14 Ferrari, S. et al. Efficient gene editing of human long-term hematopoietic stem cells validated by clonal tracking. Nat Biotechnol 38, 1298–1308, doi:10.1038/s41587-020-0551-y (2020).

15 Kuo, C. Y. et al. Site-Specific Gene Editing of Human Hematopoietic Stem Cells for X-Linked Hyper-IgM Syndrome. Cell Rep 23, 2606–2616, doi:10.1016/j.celrep.2018.04.103 (2018).

16 Pavel-Dinu, M. et al. Gene correction for SCID-X1 in long-term hematopoietic stem cells. Nat Commun 10, 1634, doi:10.1038/s41467-019-09614-y (2019).

17 Vavassori, V. et al. Modeling, optimization, and comparable efficacy of T cell and hematopoietic stem cell gene editing for treating hyper-IgM syndrome. EMBO Mol Med 13, e13545, doi:10.15252/emmm.202013545 (2021).

18 Notta, F. et al. Isolation of single human hematopoietic stem cells capable of long-term multilineage engraftment. Science 333, 218–221, doi:10.1126/science.1201219 (2011).

19 Nitsche, A. et al. Interleukin-3 promotes proliferation and differentiation of human hematopoietic stem cells but reduces their repopulation potential in NOD/SCID mice. Stem Cells 21, 236–244, doi:10.1634/stemcells.21-2-236 (2003).

20 Chaurasia, P., Gajzer, D. C., Schaniel, C., D’Souza, S. & Hoffman, R. Epigenetic reprogramming induces the expansion of cord blood stem cells. J Clin Invest 124, 2378–2395, doi:10.1172/JCI70313 (2014).

21 Hua, P. et al. Single-cell assessment of transcriptome alterations induced by Scriptaid in early differentiated human haematopoietic progenitors during ex vivo expansion. Sci Rep 9, 5300, doi:10.1038/s41598-019-41803-z (2019).

22 Milhem, M. et al. Modification of hematopoietic stem cell fate by 5aza 2’deoxycytidine and trichostatin A. Blood 103, 4102–4110, doi:10.1182/blood-2003-07-2431 (2004).

23 Zonari, E. et al. Efficient Ex Vivo Engineering and Expansion of Highly Purified Human Hematopoietic Stem and Progenitor Cell Populations for Gene Therapy. Stem Cell Reports 8, 977–990, doi:10.1016/j.stemcr.2017.02.010 (2017).

24 Biasco, L. et al. In Vivo Tracking of Human Hematopoiesis Reveals Patterns of Clonal Dynamics during Early and Steady-State Reconstitution Phases. Cell Stem Cell 19, 107–119, doi:10.1016/j.stem.2016.04.016 (2016).

25 Scala, S. et al. Dynamics of genetically engineered hematopoietic stem and progenitor cells after autologous transplantation in humans. Nat Med 24, 1683–1690, doi:10.1038/s41591-018-0195-3 (2018).

